# A screen for genes that regulate synaptic growth reveals mechanisms that stabilize synaptic strength

**DOI:** 10.1101/425876

**Authors:** Pragya Goel, Mehak Khan, Samantha Howard, Beril Kiragasi, Koto Kikuma, Dion Dickman

## Abstract

Synapses grow, prune, and remodel throughout development, experience, and disease. This structural plasticity can destabilize information transfer in the nervous system. However, neural activity remains remarkably stable throughout life, implying that adaptive countermeasures exist to stabilize neurotransmission. Aberrant synaptic structure and function has been associated with a variety of neural diseases including Fragile X syndrome, autism, and intellectual disability. We have screened disruptions in over 300 genes in *Drosophila* for defects in synaptic growth at the neuromuscular junction. This effort identified 12 mutants with severe reductions or enhancements in synaptic growth. Remarkably, electrophysiological recordings revealed synaptic strength in all but one of these mutants was unchanged compared to wild type. We utilized a combination of genetic, anatomical, and electrophysiological analyses to illuminate three mechanisms that stabilize synaptic strength in the face of alterations in synaptic growth. These include compensatory changes in 1) postsynaptic receptor abundance; 2) presynaptic morphology; and 3) active zone structure. Together, this analysis identifies new genes that regulate synaptic growth and the adaptive strategies that synapses employ to homeostatically stabilize synaptic strength in response.

**AUTHOR SUMMARY:** Throughout development, maturation, experience, and disease, synapses undergo dramatic changes in growth and remodeling. Although these processes are necessary for learning and memory, they pose major challenges to stable function in the nervous system. However, neurotransmission is typically constrained within narrow physiological ranges, implying the existence of homeostatic mechanisms that maintain stable functionality despite drastic alterations in synapse number. In this study we investigate the relationship between synaptic growth and function across a variety of mutations in neural and synaptic genes in the fruitfly *Drosophila melanogaster*. Using the neuromuscular junction as a model system, we reveal three adaptive mechanisms that stabilize synaptic strength when synapses are dramatically under- or over-grown. Together, these findings provide insights into the strategies employed at both pre- and post-synaptic compartments to ensure stable functionality while allowing considerable flexibility in overall synapse number.

## INTRODUCTION

Dramatic changes in synapse number, morphology, and structure occur throughout nervous system development and during various forms of plasticity and remodeling in the mature nervous system. For example, expansion and retraction of synaptic terminals contributes to the refinement of neural circuits during developmental pruning, sleep/wake behavior, and experience-dependent plasticity [1-4]. While these dynamic changes enable the flexibility necessary to wire the nervous system during development and to modify synapses during learning and memory, they pose a major challenge to the stability of neural function. Indeed, it is interesting to note that the period of highest susceptibility to seizures occurs during the first years of life, a period of dramatic growth and pliability in the brain [5, 6]. However, despite the potential for these processes to disrupt information transfer in the nervous system, homeostatic mechanisms maintain physiologically stable levels of functionality [7, 8]. Although the genes and molecular processes that enable synapse-specific control of Hebbian and homeostatic plasticity have been intensively studied [9-11], how global levels of synaptic strength are stabilized and integrated with local mechanisms remains enigmatic.

The *Drosophila* neuromuscular junction (NMJ) is a powerful model system to illuminate the genes and mechanisms that regulate synaptic growth, function, and homeostatic plasticity. At this model glutamatergic synapse, stereotyped levels of synaptic strength are consistently observed despite a dramatic expansion of synaptic growth, where the NMJ rapidly enlarges by ~100-fold during larval development [12, 13]. Remarkably, synaptic strength is maintained within narrow physiological ranges during this process [14], implying that homeostatic processes stabilize neurotransmission in coordination with synaptic growth. A variety of homeostatic mechanisms are triggered at the *Drosophila* NMJ in response to excess glutamate release [15-17], diminished postsynaptic neurotransmitter receptor functionality [18, 19], injury-related signaling [20], and biased innervation [20, 21]. These mechanisms can operate with specificity at a subset of synapses [21-23]. However, there is evidence that additional homeostatic processes stabilize global synaptic strength when total synapse numbers are drastically altered at the NMJ. For example, it has been estimated that as many as 44% of the genes encoded in the *Drosophila* genome influence synaptic growth and structure [24], while far fewer genes appear to be involved in neurotransmission [25, 26]. Despite these observations, the mechanisms that stabilize global synaptic strength in the face of variations in synaptic growth have yet to be defined.

Genes that have been linked to neurological and neuropsychiatric diseases are attractive candidates screen for roles in regulating synaptic growth, structure, and plasticity. Aberrant synaptic growth, structure, and plasticity is associated with a variety of neural diseases including Fragile X Syndrome, autism spectrum disorder, schizophrenia, and intellectual disability [27, 28]. For example, the Fragile X Mental Retardation protein (FMRP), an RNA binding protein, modulates translation and targets hundreds of synaptic genes in both pre- and post-synaptic compartments to sculpt synaptic structure and function [29-32]. Recent biochemical and next-generation sequencing approaches have identified over 800 transcripts that associate with FMRP [33, 34]. Further, emerging genetic linkage studies have implicated a variety of synaptic genes associated with susceptibility to autism, schizophrenia, bipolar disorder, and intellectual disability [35-37]. Hence, screening genes linked with neural diseases provides a compelling foundation to define new genes with fundamental roles at synapses.

We have systematically screened a collection of genes with links to neural diseases for roles in synaptic growth and transmission at the *Drosophila* NMJ. This analysis discovered several new genes required for proper synaptic growth and transmission. Interestingly, this approach also confirmed that while synaptic growth can vary considerably across mutations in diverse genes, neurotransmission is constrained within much narrower physiological ranges. Given these results, we chose not to characterize in detail the specific functions of individual genes in regulating synaptic growth. Rather, we investigated synaptic structure and function in the subset of mutants that exhibited the most extreme changes in synaptic growth but that, remarkably, maintained stable synaptic strength. This effort defined three mechanisms that targeted both pre- and post-synaptic structures for homeostatic modulation. Together, these results elucidate adaptive strategies that can be employed by synapses to maintain set point levels of synaptic strength when confronted with extreme alterations to synaptic growth.

## RESULTS

### A forward genetic screen identifies genes that regulate synaptic growth and transmission at the *Drosophila* NMJ

To systematically screen a collection of genes for roles in synaptic growth and function, we first established a list of *Drosophila* homologs of mammalian genes linked to synaptic function and neural disease. The initial list consisted of ~800 mammalian genes expressed at synapses and/or linked with neural disease (S1 Table). These genes included putative transcripts associated with FMRP [34, 38] and additional genes that have been associated with schizophrenia and autism spectrum disorder [39-42]. From this list, we identified a final group of 300 *Drosophila* homologues - 132 putative FMRP targets and 168 genes associated with synapses or other diseases. From this initial list, we obtained a collection of 109 putative genetic mutations and 191 RNAi lines from public resources (S1 Table). Finally, we assessed the lethal phase of homozygous mutants and RNAi lines crossed to NMJ drivers, removing any that failed to survive to at least the third-instar larval stage. Together, this effort established a collection of 297 stocks to screen for defects in synaptic growth and function at the third-instar larval NMJ.

We first assessed synaptic growth in this collection of 297 mutants and RNAi lines. Specifically, we characterized homozygous mutants or larvae in which RNAi transgenes were driven in both motor neurons and muscle (see Methods; [43]). Immunostaining of synaptic boutons at the *Drosophila* NMJ was used to quantify synaptic growth. Wild-type NMJs typically exhibit ~30 boutons at the muscle 4 NMJ (Fig 1A and 1B and 1D). We immunostained the NMJ with a markers for synaptic vesicles (vGlut) and the neuronal membrane (HRP), and considered a single puncta of vGlut intensity to represent a synaptic bouton (Fig 1A and 1B). Quantification of bouton numbers across all 297 mutants and RNAi lines revealed a broad distribution, with 31.2 boutons as the mean and a standard deviation of 6.8 (Fig 1D). From this analysis, we selected the subset of mutants or RNAi lines that displayed the most extreme difference in bouton number, using two standard deviations above or below the mean (> 44% increase or decrease; Fig 1C and 1D) as cutoffs for further study.

**Fig 1.**
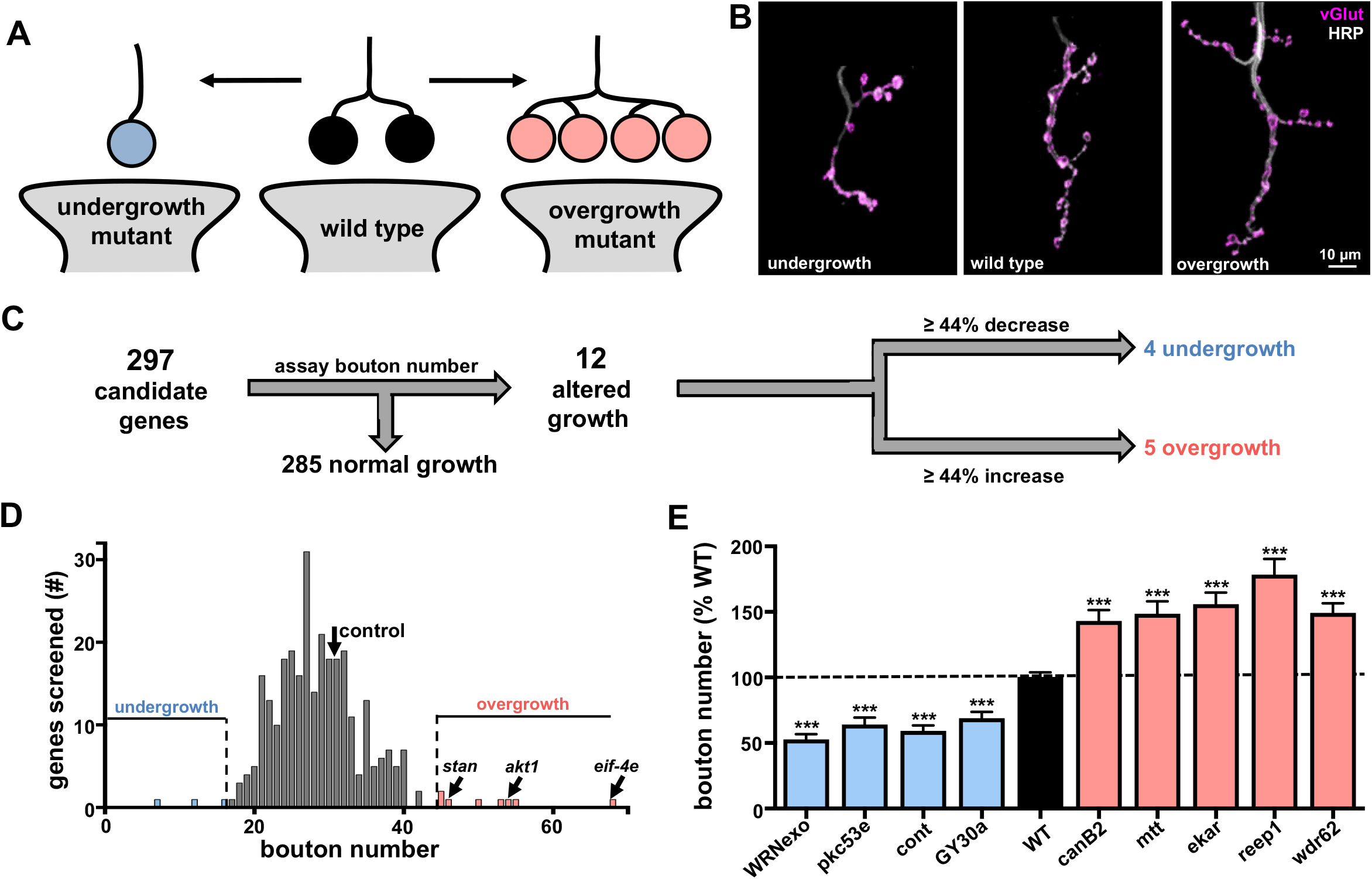
A forward genetic screen identifies genes that regulate synaptic growth at the *Drosophila* NMJ. **(A)** Schematic illustrating synaptic boutons, numbers of which are a measure of NMJ growth. **(B)** Images of larval muscle 4 NMJs immunostained with anti-HRP (neuronal membrane marker) and anti-vGlut (synaptic vesicle marker). Examples of NMJs in undergrowth and overgrowth mutants are shown. **(C)** Flow diagram of synaptic growth screen strategy and outcome. Mutants with increases or decreases in synaptic growth that were over 2 standard deviations from controls (~44% increase or decrease) are indicated. **(D)** Histogram of average bouton number per mutant or RNAi line quantified in the synaptic growth screen. Average bouton numbers in control (black arrow), overgrowth mutants (red), and undergrowth mutants (blue) are indicated. Three genes previously reported to exhibit synaptic overgrowth are indicated. **(E)** Bouton numbers of the identified overgrowth and undergrowth mutants shown as a percentage of wild-type values. No significant differences in bouton numbers were observed between the mutant control (*w^1118^*) and RNAi line control (*C15xUAS-RFP*; S1 Table), so all values were pooled. Error bars indicate ±SEM. ***p≤0.001. Additional details of all mutants and RNAi lines screened and statistical information (mean values, SEM, n, p) are shown in S1 Table.

12 targets with extreme changes in synaptic growth at the NMJ were identified (Fig 1C and 1D). All 12 were genetic mutants; four exhibited a reduction of over 44% in bouton number and were termed “undergrowth mutants” (Fig 1C-1E; blue), while the other eight exhibited an increase of over 44% in bouton number and were termed “overgrowth mutants” (Fig 1C-1E; red). Of the 12 positive hits from our initial screen, three genes were previously reported to have defects in synaptic growth (Fig 1D and 1E), serving to validate our approach. These include the G-protein-coupled receptor *flamingo* [44], the serine-threonine kinase *Akt1* [45], and the translation factor *eIF-4E* [46-48]. Thus, from this initial screen of 297 lines, we identified four undergrowth and five overgrowth genes, which have not previously been reported to regulate synaptic growth. The putative functions of these genes are detailed in S2 Table.

We also assayed synaptic transmission in the collection of 297 lines. We used electrophysiology to quantify miniature excitatory postsynaptic potential (mEPSP) amplitude, evoked excitatory postsynaptic potential (EPSP) amplitude, and to calculate the number of synaptic vesicles released per stimulus (quantal content, a measure of neurotransmitter release) from each mutant screened (S1 Table). Electrophysiological recordings from all mutant and RNAi lines revealed a mean EPSP amplitude of 35.4 mV and a standard deviation of 6.5 mV (Fig 2B). We identified 40 mutant and RNAi lines with EPSP amplitudes over two standard deviations below the mean (>36%; Fig 2A and 2B), while no targets exhibited an increase in EPSP amplitude of >36% relative to the mean (Fig 2A; S1 Table). Quantification of bouton numbers in the 40 synaptic transmission mutants or RNAi lines revealed values similar to wild type (Fig 2C), consistent with previous studies that have shown aberrant synaptic function often occurs without any major defects in synaptic growth [25, 49, 50]. This suggests defects in synaptic function alone, independently of reduced growth, disrupts synaptic strength in these lines.

**Fig 2.**
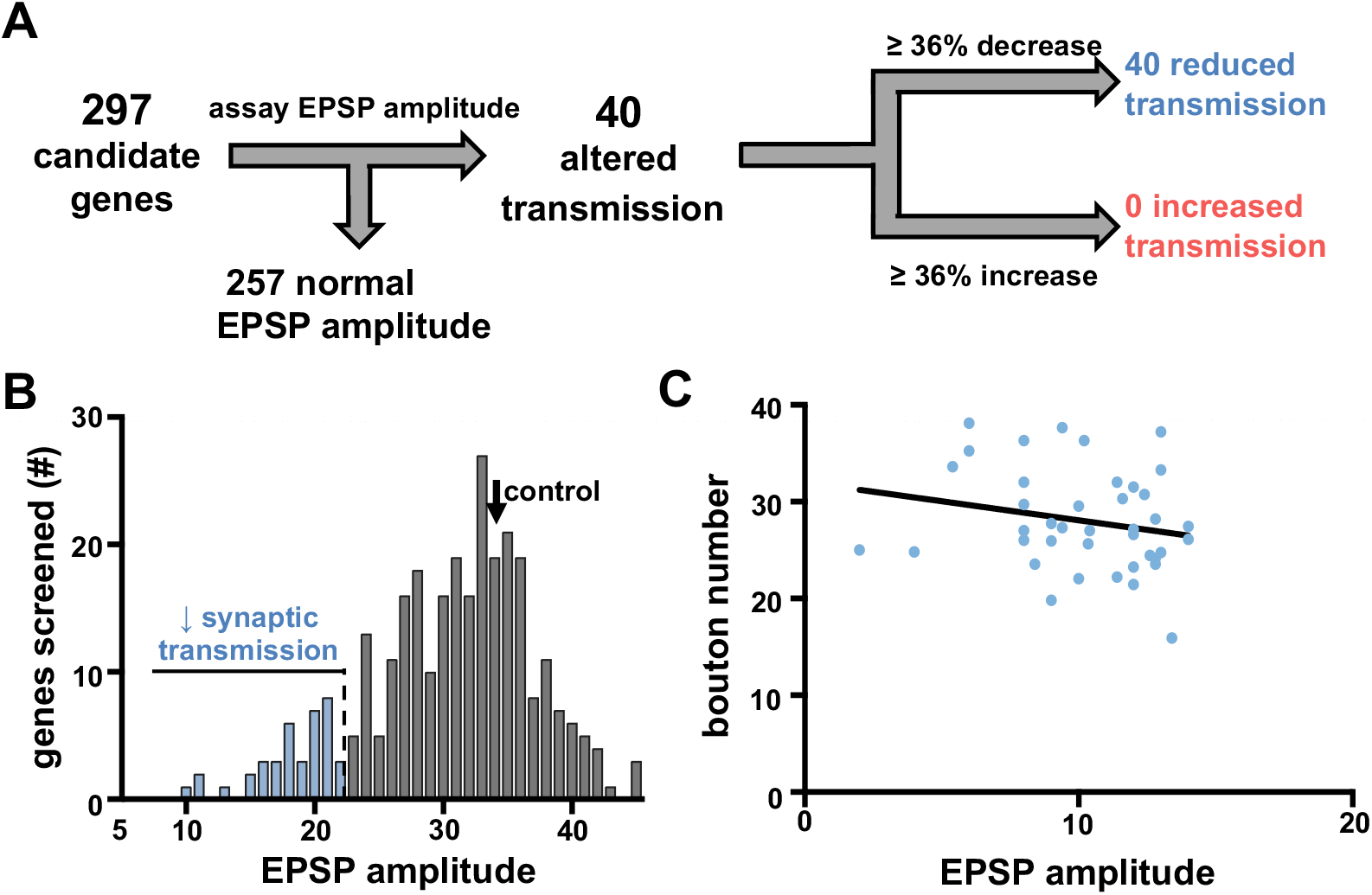
Presynaptic neurotransmitter release does not scale with synaptic growth in the mutants screened. **(A)** Flow diagram of electrophysiology-based synaptic transmission screen strategy and outcome. **(B)** Histogram of average EPSP amplitude quantified for each mutant and RNAi line in the screen. Although no mutants or RNAi lines with EPSP amplitudes > two 22 standard deviations above the average EPSP amplitude in wild type were found (~36% increase), several lines with EPSP amplitudes below this threshold were dentified (indicated in blue). **(C)** Graph showing the total bouton number of each synaptic transmission mutant or RNAi line identified as a function of EPSP amplitude. The best-fit line to this data (solid black line; slope = -0.788) indicates that bouton numbers do not correlate with EPSP amplitude (R^2^= 0.045, p value=0.186). Additional details of all mutant and RNAi lines screened and statistical information (mean values, SEM, n, p) are shown in S1 Table.

### Synaptic strength remains constant despite variations in synaptic growth

We focused on understanding how synaptic function remains stable across the broad variation in synaptic growth by analyzing synaptic growth and structure in the 257 remaining mutants and RNAi lines with relatively stable EPSP amplitudes. First, we considered two possible models to describe the relationship between synaptic growth (bouton numbers) and synaptic strength (EPSP amplitude). In a “scaling” model, each individual bouton functions as an independent unit of synaptic function, with all boutons functionally equivalent (Fig 3A). Hence, synaptic strength would be predicted to scale in amplitude in proportion to the total number of synaptic boutons, with the number of individual synapses (active zone and glutamate receptor dyads) linearly increasing with the number of boutons. Assuming the functionality of each dyad to be constant, as bouton number increases or decreases, total synaptic strength would scale accordingly (Fig 3A). Alternatively, in a “homeostatic” model, synapses would be adaptively modulated to counteract variations in synaptic growth and maintain stable levels of global synaptic strength (Fig 3B). In this case, adaptations in total active zone number, presynaptic release probability, and/or postsynaptic receptivity to neurotransmitter would compensate for altered bouton number to tune synaptic strength and maintain constant levels of neurotransmission. We considered whether a scaling or homeostatic model best described our data from the genetic screen.

**Fig 3.**
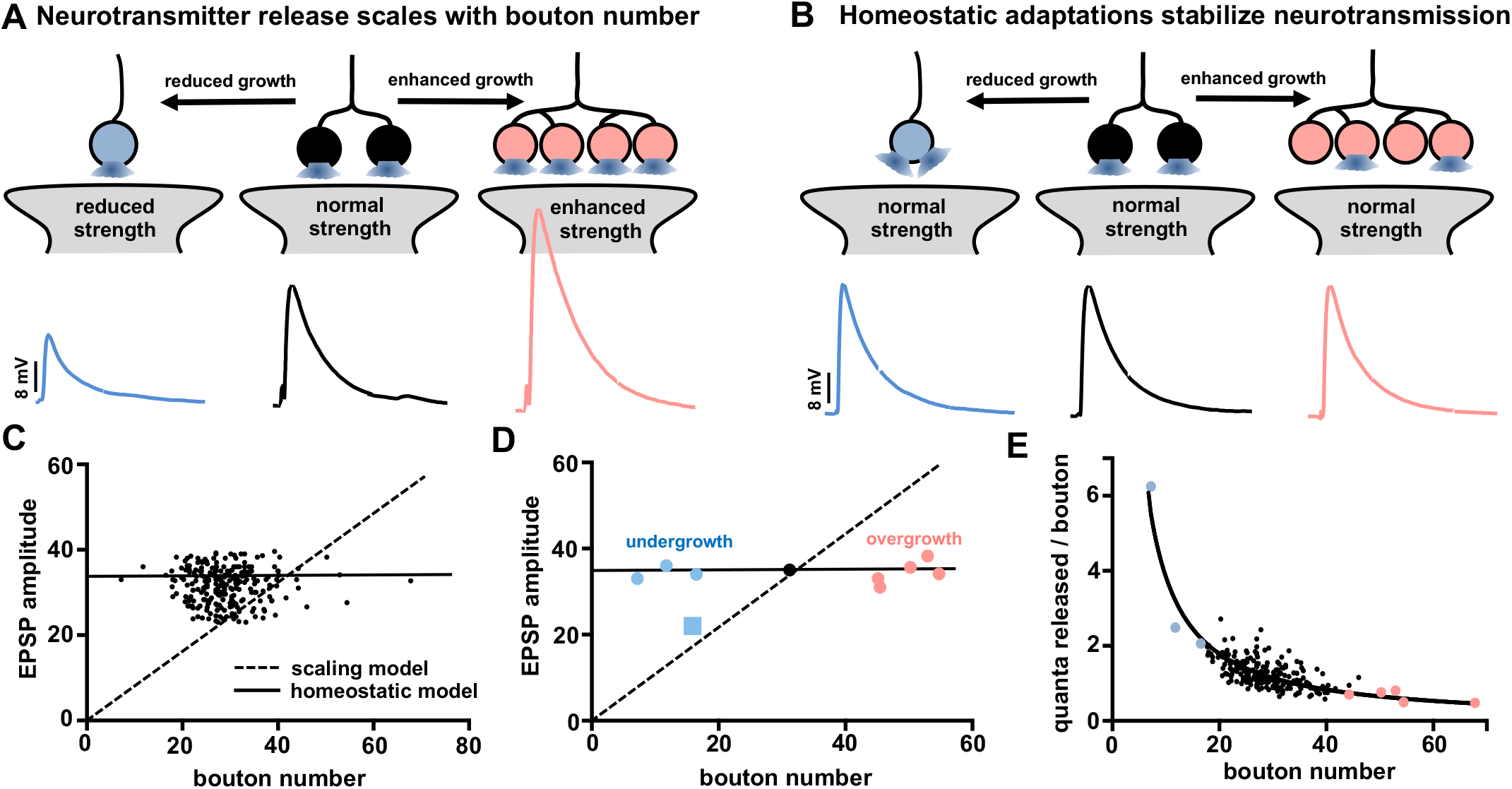
Stable synaptic strength is observed despite variation in synaptic growth in the mutants screened. **(A)** Schematic illustrating a “scaling” model in which presynaptic neurotransmitter release scales with synaptic growth. Note that in this scenario, EPSP amplitude correlates with bouton number. **(B)** Schematic illustrating an alternative “homeostatic” model, in which synaptic strength remains constant across changes in bouton number. **(C)** Graph plotting the EPSP amplitude of the genes screened (with the mutants and RNAi lines defective in synaptic transmission removed) plotted as a function of bouton number. The dashed diagonal line represents the ideal “scaling” model, where EPSP amplitude correlates with bouton numbers. The horizontal solid line represents the idealized “homeostatic” model, where no such correlation is observed. The data shows that EPSP amplitudes do not correlate with bouton numbers (Pearson’s correlation coefficient R^2^= 0.0002, p value=0.789), a closer fit to a “homeostatic” model. **(D)** Graph plotting EPSP amplitude of the synaptic overgrowth and undergrowth mutants as a function of bouton number. Only a single undergrowth mutant (indicated as a square data point) fits the “scaling” model, with EPSP amplitude reduced to a similar extent as the reduction in bouton number. All other synaptic growth mutants maintained stable EPSP amplitude, consistent with a “homeostatic” model (solid horizontal line; Pearson’s correlation coefficient R^2^= 0.012, p value=0.718). **(E)** Average quanta released per bouton calculated for each mutant is plotted as a function of bouton number for the mutants shown in (C). A curve fit of this data provides a Goodness of Fit R^2^ value of 0.65 and a p value of < 0.0001, indicating an inverse correlation between quanta released per bouton with total bouton number. Additional details of the mutants screened and statistical information (mean values, SEM, n, p) are shown in S1 Table.

We plotted the average EPSP amplitude of each mutant screened as a function of total bouton number for that specific mutant (Fig 3C). A scaling model would predict a linear relationship in this plot, where synaptic strength (EPSP amplitude) is proportional to bouton number (indicated by the dotted line in Fig 3C). However, this analysis found no significant correlation between EPSP amplitude and bouton number (R^2^=0.0002, p-value=0.7935). Rather, the majority of mutants screened (86%) maintained EPSP amplitudes of 32-36 mV (Fig 3C), more consistent with a homeostatic model. Next, we examined synaptic strength in the most extreme four undergrowth and five overgrowth mutants discussed in Fig 1E. We plotted the EPSP amplitude for each mutant as a function of bouton number (Fig 3D). Interestingly, all but one of the nine mutants exhibited EPSP amplitudes consistent with a homeostatic model, while one mutant, *pkc53E*, best fit with a scaling model. Finally, we considered that for a homeostatic model to be truly “homeostatic”, presynaptic neurotransmitter release (quantal content) for each individual bouton should inversely scale with total boutons per NMJ. Indeed, when the average quantal content was normalized per bouton for all 257 mutants and RNAi lines, a robust scaling of quanta released per bouton was observed (Fig 3E), consistent with a homeostatic tuning of presynaptic release per bouton. Together, this analysis of synaptic growth and function in the genes screened is consistent with the homeostatic model schematized in Fig 3B, suggesting that presynaptic release is tuned at individual boutons to maintain stable global synaptic strength despite variation in synaptic growth.

### Synaptic strength scales with synaptic growth in *pkc53E* mutants

We next sought to characterize the relationship between synaptic growth and function in the nine FMRP target mutants in more detail. In particular, we sought to illuminate how, or whether, synaptic scaling or homeostasis was expressed. We first characterized synaptic function and structure in the four undergrowth mutants. Mutations in the first gene, *protein kinase C 53E* (*pkc53E*), exhibited reductions in synaptic strength that appeared to scale with synaptic growth (Fig 3D). Bouton numbers were reduced by ~50% in homozygous mutants of *pkc53E* (S1 Table) and in *pkc53E* mutants in trans to a deficiency that removed the entire locus (*pkc53E^1^/pkc53E^Df^*; Fig 4A and 4B; S3 Table). Correspondingly, EPSP amplitude was reduced to a similar extent in both allelic combinations of *pkc53E* compared to wild type (Fig 4C and 4D). Synaptic strength, indicated by EPSP amplitude, is determined by two parameters: The amount of presynaptic neurotransmitter released and the postsynaptic response to neurotransmitter [51, 52]. A change in mEPSP amplitude, which reflects the postsynaptic response to neurotransmitter released from a single vesicle, would likely indicate a change in the number or functionality of postsynaptic glutamate receptors in *pkc53E* mutants. However, we observed no significant difference in mEPSP amplitude in *pkc53E* mutants compared to wild type (Fig 4D; S3 Table), consistent with no postsynaptic adaptations in this mutant. Next, we calculated quantal content in these mutants; a measure of the number of synaptic vesicles released in response to synaptic stimulation, and found a reduction in this value proportional to the reduction in EPSP amplitude (Fig 4C and 4D), as expected. If no adaptions to presynaptic structure occurred in *pkc53E*, then the anatomical number of release sites (active zones) should be reduced in proportion to the reduction in bouton number. We measured the number of puncta of the active zone scaffold Bruchpilot (BRP) by immunostaining the NMJ, which represent individual releases sites [49, 53]. We observed a reduction in BRP puncta number per NMJ proportional to the reduction in bouton number in *pkc53E* mutants (Fig 4E and 4F), with no change in BRP puncta density compared to wild type (Fig 4G). Thus, in *pkc53E* mutants, the number of active zones is reduced in proportion to the number of boutons and no apparent changes are observed in release probability or the postsynaptic sensitivity to neurotransmitter, consistent with a scaling of synaptic strength with synaptic growth. Importantly, this implies that in the remaining eight mutants in which synaptic strength remained constant despite increased or reduced growth, some compensatory adaptions must have occurred.

**Fig 4.**
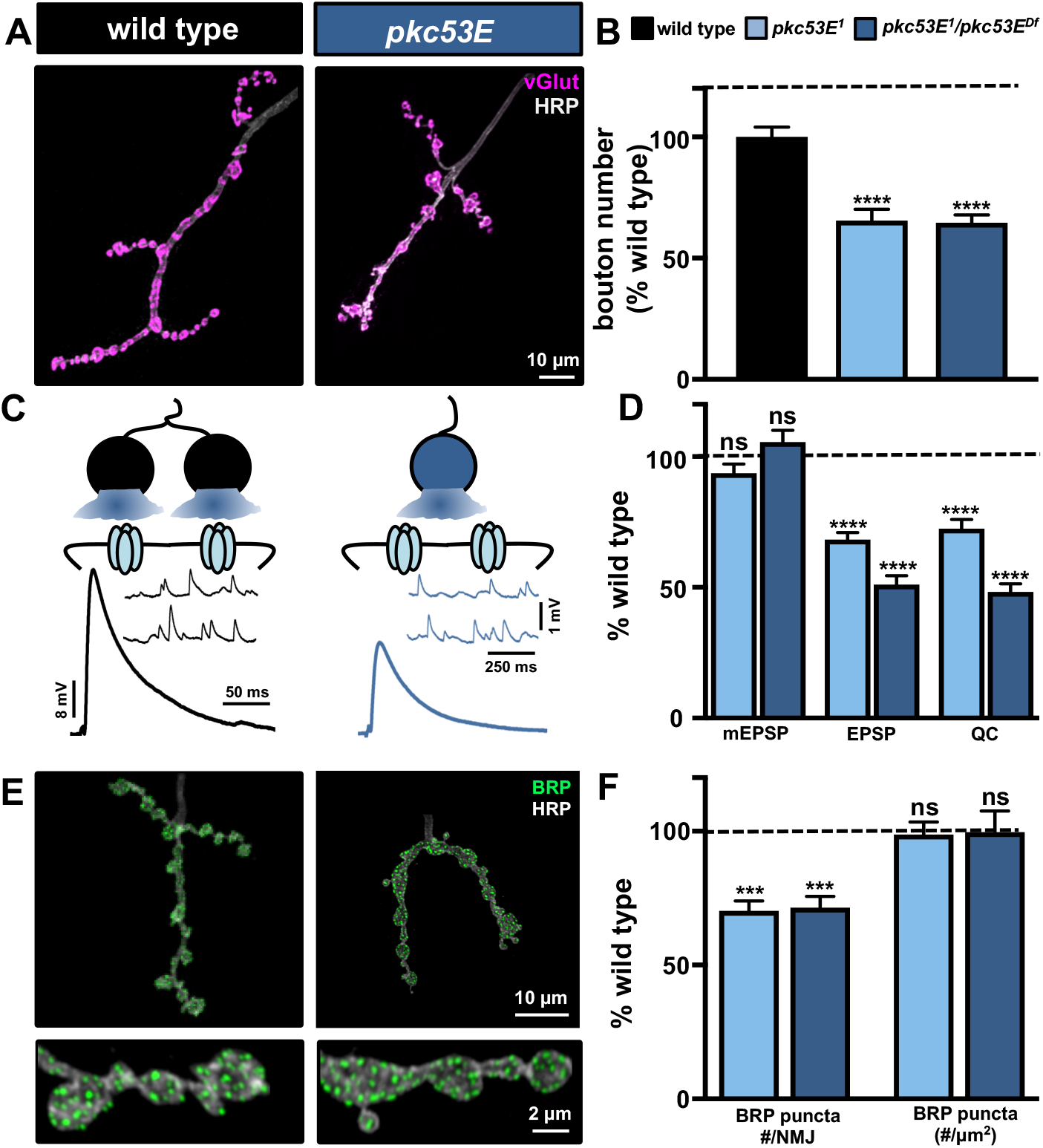
Presynaptic neurotransmitter release scales with reduced bouton and active zone number in *pkc53E* mutants. **(A)** Representative muscle 4 NMJ images of wild type (*w^1118^*) and *pkc53E* mutants *in trans* with a deficiency (*pkc53E^1^/pkc53E^Df(2R)P803-Delta15^*) immunostained with anti-HRP and anti-vGlut. **(B)** Quantification of bouton number in the indicated genotypes normalized to wild-type values. **(C)** Schematic and representative electrophysiological traces of mEPSPs and EPSPs in the indicated genotypes illustrating reduced synaptic strength and no evidence for compensatory adaptions to presynaptic neurotransmitter release or postsynaptic sensitivity to neurotransmitter. **(D)** Quantification of mEPSP, EPSP, and quantal content values in *pkc53E* mutants normalized as a percentage of wild type. **(E)** Representative images of NMJs immunostained with anti-HRP and the anti-bruchpilot (BRP; presynaptic active zone marker), with individual boutons shown at higher magnification (insets below). **(F)** Quantification of total BRP puncta number per NMJ shows a concomitant reduction with bouton number and no significant change in BRP puncta density. Error bars indicate ±SEM. One-way analysis of variance (ANOVA) test was performed, followed by a Tukey’s multiple-comparison test. ***p≤0.001; ****p≤0.0001; ns=not significant, p>0.05. Detailed statistical information (mean values, SEM, n, p) is shown in S3 Table.

### Enhanced postsynaptic receptor abundance compensates for reduced presynaptic release in *WRNexo* mutants

We next focused on the undergrowth mutant *WRNexo*. *WRNexo* encodes *Werner’s exonuclease*, so named because mutations in the human homolog cause the disease Werner’s Syndrome, a disease resulting in premature aging due to DNA damage [54-56]. Null mutations in *WRNexo* have been generated and characterized in the context of DNA repair in *Drosophila* [57]. However, roles for *WRNexo* in synaptic growth or function have not been reported, nor have they been characterized at the NMJ. *WRNexo* mutants exhibit significant reductions in synaptic growth, with bouton numbers reduced by ~50% compared to wild type controls (Fig 5A and 5B). However, EPSP amplitude in *WRNexo* mutants was similar to wild type (Fig 5C and 5D). Quantification of mEPSP amplitude revealed a significant increase in *WRNexo* mutants compared to wild type, resulting in a corresponding reduction in quantal content (Fig 5C and 5D). Together, this suggests that while presynaptic neurotransmitter release is reduced in accordance to reduced synaptic growth in *WRNexo* mutants, an increase in the postsynaptic responsiveness to neurotransmitter was sufficient to maintain normal synaptic strength.

**Fig 5.**
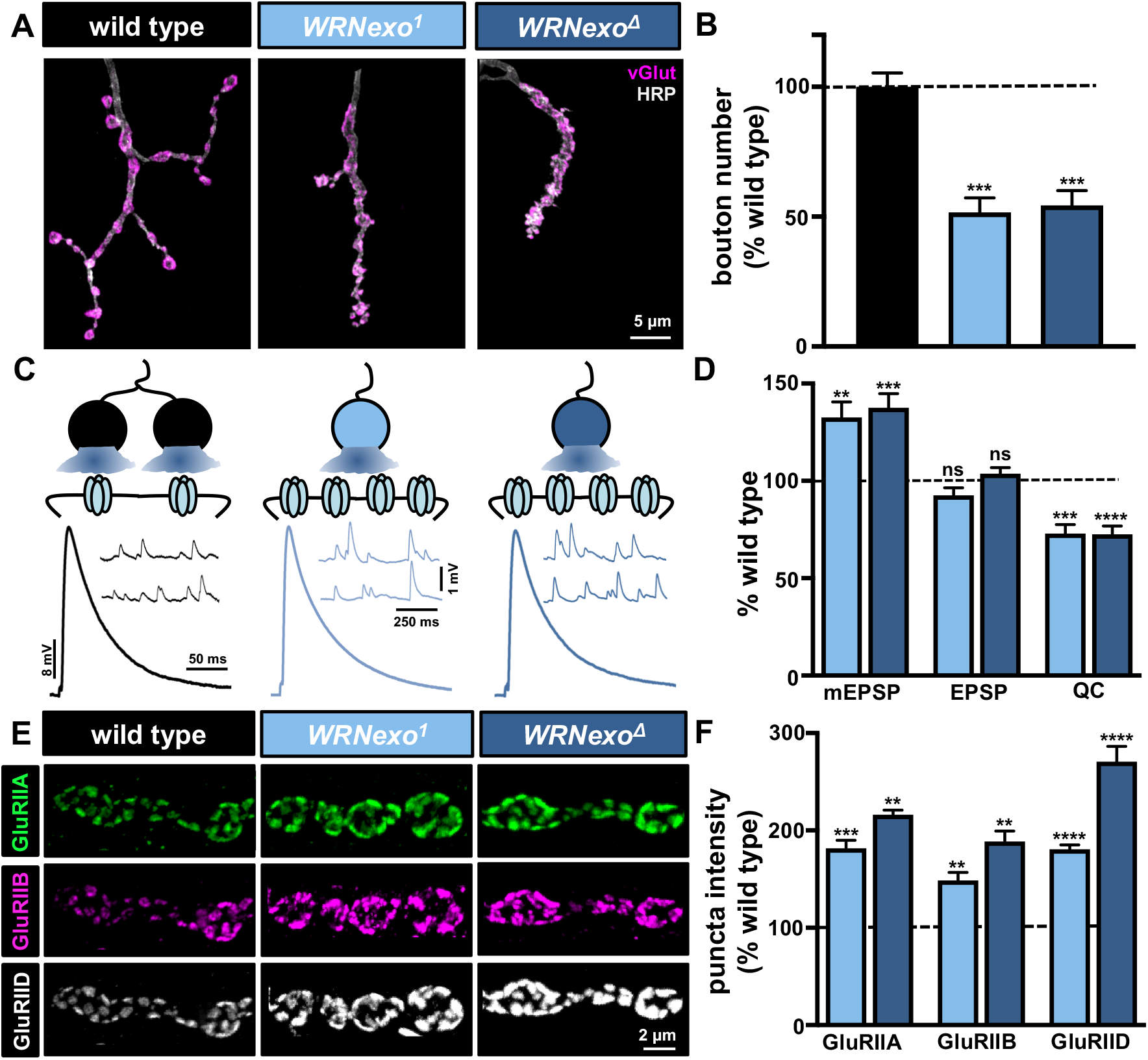
Increased postsynaptic receptor levels compensate for reduced presynaptic neurotransmitter release in *WRNexo* mutants. **(A)** Representative images of muscle 4 NMJs in wild type, *WRNexo* mutants (*WRNexo^MI13095^*), and *WRNexo* null mutants (*WRNexo*^D^), immunostained with anti-HRP and anti-vGlut. **(B)** Quantification of bouton numbers in *WRNexo* mutants normalized as a percentage of wild type. **(C)** Representative mEPSP and EPSP traces in the indicated genotypes. The schematic illustrates that enhanced levels of postsynaptic glutamate receptor levels offset reduced presynaptic release in *WRNexo* mutants. **(D)** Quantification of mEPSP, EPSP, and quantal content values in the indicated genotypes normalized as a percentage of wild type. **(E)** Representative images of boutons immunostained with antibodies against three postsynaptic glutamate receptor subunits (GluRIIA; GluRIIB; GluRIID). **(F)** Quantification of sum puncta fluorescence intensity of each receptor subunit reveals enhanced levels of all postsynaptic receptors in *WRNexo*. Error bars indicate ±SEM. One-way analysis of variance (ANOVA) test was performed, followed by a Tukey’s multiple-comparison test. **p≤0.01; ***p≤0.001; ****p≤0.0001; ns=not significant, p>0.05. Detailed statistical information (mean values, SEM, n, p) is shown in S3 Table.

At the *Drosophila* NMJ, two glutamate receptor subtypes, GluRIIA-containing and GluRIIB-containing, mediate the response to synaptically released glutamate [58]. Three essential glutamate receptors, GluRIIC, GluRIID, and GluRIIE are core components of both receptor complexes and incorporate either GluRIIA or GluRIIB subunits [58, 59]. The majority of neurotransmission is driven by GluRIIA-containing receptors due to their slower desensitization kinetics and larger current amplitudes [19, 60, 61]. Given the increase in mEPSP amplitude observed in *WRNexo* mutants, we examined the state of glutamate receptors in more detail. We co-stained NMJs with antibodies against GluRIIA, GluRIIB, and GluRIID and assessed the synaptic localization of these receptor subunits while also quantifying immunofluorescence levels (Fig 5E and 5F). While we did not observe any major differences in the localization of receptors at the NMJ, we did find a significant increase in GluRIIA, GluRIIB and GluRIID subunit levels in *WRNexo* mutants (Fig 5E and 5F). This suggests that the additional abundance of postsynaptic glutamate receptors at the postsynaptic density of *WRNexo* mutants increased sensitivity to glutamate and compensated for reduced synaptic growth and glutamate release.

### Increased bouton area maintains stable synapse number in *cont* and *Gγ30a* mutants

Next, we characterized the two remaining synaptic undergrowth mutants, *contactin* (*cont*), a cell adhesion molecule involved in septate junction organization between glia and neurons [62], and *G γ30A*, the gamma subunit of a heterotrimeric G protein [63]. Interestingly, despite a ~60% reduction in bouton number compared to wild type (Fig 6A and 6B), these two mutants appeared to have no obvious changes in synaptic physiology (Fig 6C and 6D). mEPSP amplitudes were similar to wild type in both mutants, which implies that a presynaptic change in either active zone number and/or release probability likely compensated for reduced bouton number to maintain stable levels of presynaptic neurotransmitter release.

**Fig 6.**
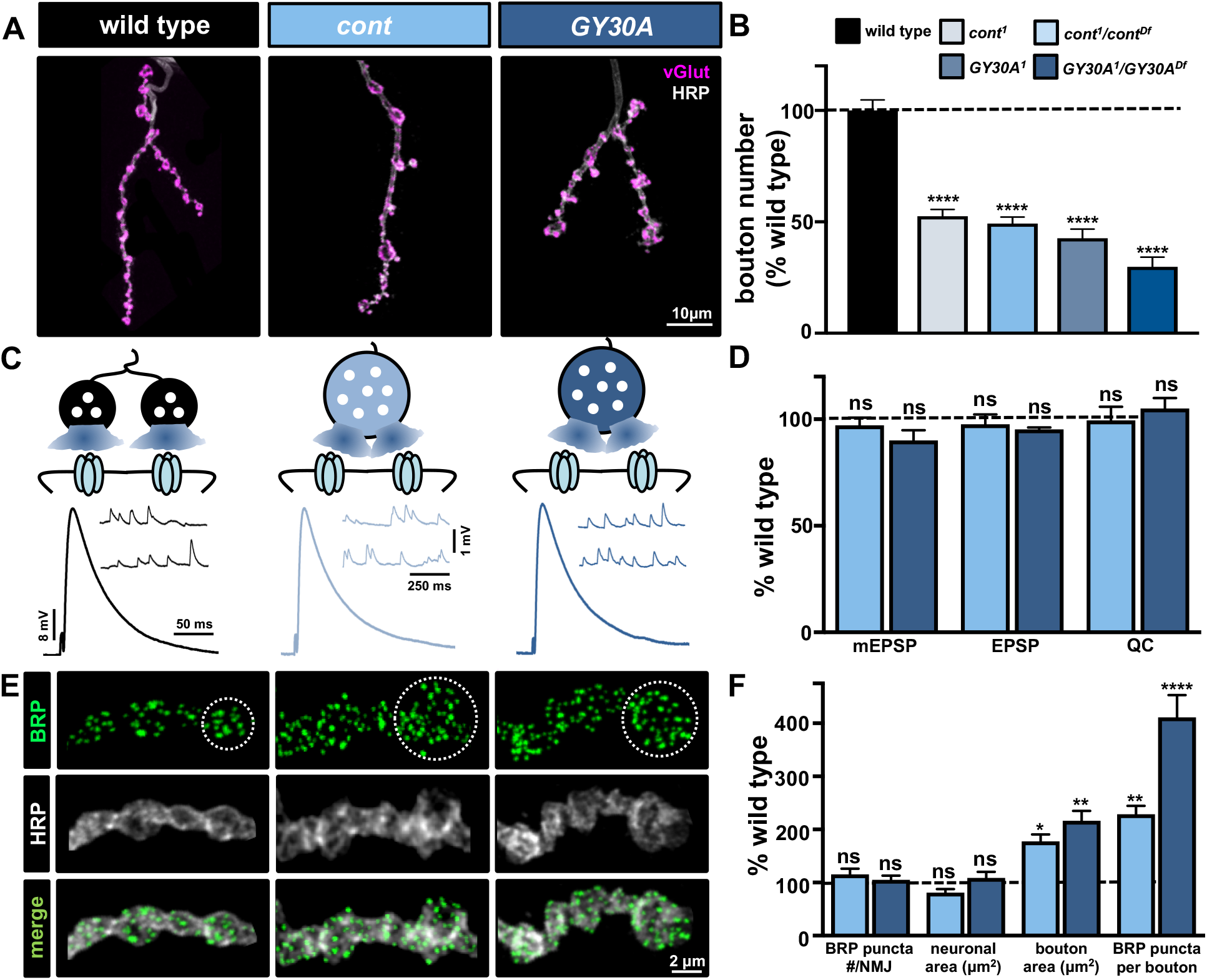
Increased bouton size compensates for reduced bouton number in *cont* and *G*γ*30A* mutants. **(A)** Representative images of muscle 4 NMJs in wild type, *cont* and *Gγ30A* mutants *in trans* with deficiencies (*cont*: *cont^1^*/*cont^Df(3R)BSC146^* and *Gγ30A: Gγ30A^1^*/*Gγ30A^Df(2L)ED680^*) immunostained with anti-HRP and anti-vGlut. **(B)** Bouton numbers per NMJ in the indicated genotypes normalized as a percentage of wild-type values. **(C)** Representative mEPSP and EPSP traces in the indicated genotypes. The schematic illustrates an enhancement in bouton area resulting in more release sites per bouton, with no apparent change in postsynaptic sensitivity to glutamate in *cont* and *Gγ30A* mutants. **(D)** Quantification of mEPSP, EPSP, and quantal content values in the indicated genotypes normalized as a percentage of wild type values. **(E)** Representative images of individual boutons from the indicated genotypes immunostained with anti-BRP and anti-HRP. The white circle outlines a single bouton. The increased area of individual boutons and number of BRP puncta within each bouton is apparent in *cont* and *Gγ30A* mutants. **(F)** Quantification of the indicated synaptic parameters in the indicated genotypes normalized as a percentage of wild-type values. Note that total neuronal membrane area is unchanged in *cont* and *Gγ30A* mutants due to an increase in the average area of individual boutons. Hence, a significant increase in the number of BRP puncta per bouton is observed. Error bars indicate ±SEM. One-way analysis of variance (ANOVA) test was performed, followed by a Tukey’s multiple-comparison test. *p≤0.05; **p≤0.01; ****p≤0.0001; ns=not significant, p>0.05. Detailed statistical information (mean values, SEM, n, p) is shown in S3 Table.

We therefore quantified the number of BRP puncta per NMJ in *cont* and *Gγ30A* mutants. Surprisingly, immunostaining of BRP revealed that total puncta number per NMJ were similar in both *cont* and *Gγ30A* mutants to wild type (Fig 6E and 6F). Further analysis found that while bouton numbers were indeed reduced, individual boutons were significantly enlarged in area in these mutants (Fig 6E and 6F). Thus, although *cont* and *Gγ30A* were defined as synaptic undergrowth mutants based on our bouton counting assay, increased bouton area conserved total neuronal membrane area (Fig 6F). Consistently, quantification of BRP puncta per bouton revealed a significant increase in both *cont* and *Gγ30A* (Fig 6E and 6F), demonstrating that active zone number scaled with the enhanced NMJ membrane and area of individual boutons. Thus, despite a reduction in overall bouton number, increased synapse number per bouton was sufficient to maintain total synapse number per NMJ, and synaptic strength, in both *cont* and *Gγ30A* undergrowth mutants.

### Reduced active zone area is observed in overgrowth mutants with increased active zone numbers

We next characterized synaptic function and structure in the five synaptic overgrowth mutants. This category harbored mutations in diverse genes encoding the G-protein coupled receptor *mangetout* (*mtt*); the *WD repeat domain protein 62* (*wdr62*); the kainate receptor *ekar*; the calcium-activated protein phosphatase *calcineurin B2* (*canB2*); and the endoplasmic reticulum stress gene *receptor expression enhancing protein* (*reep*). Despite the diverse functions of these genes (S2 Table), they shared a common 40-50% increase in the number of synaptic boutons per NMJ but stable synaptic strength (Fig 7A and 7C). Electrophysiological analysis revealed no significant changes in mEPSP amplitude, EPSP amplitude, or quantal content (Fig 7B and 7E; S1 Table). This suggests the postsynaptic sensitivity to neurotransmitter was not impacted in these mutants, and implies a change in synapse number and/or release probability likely compensated for the increased bouton number shared in these mutants.

**Fig 7.**
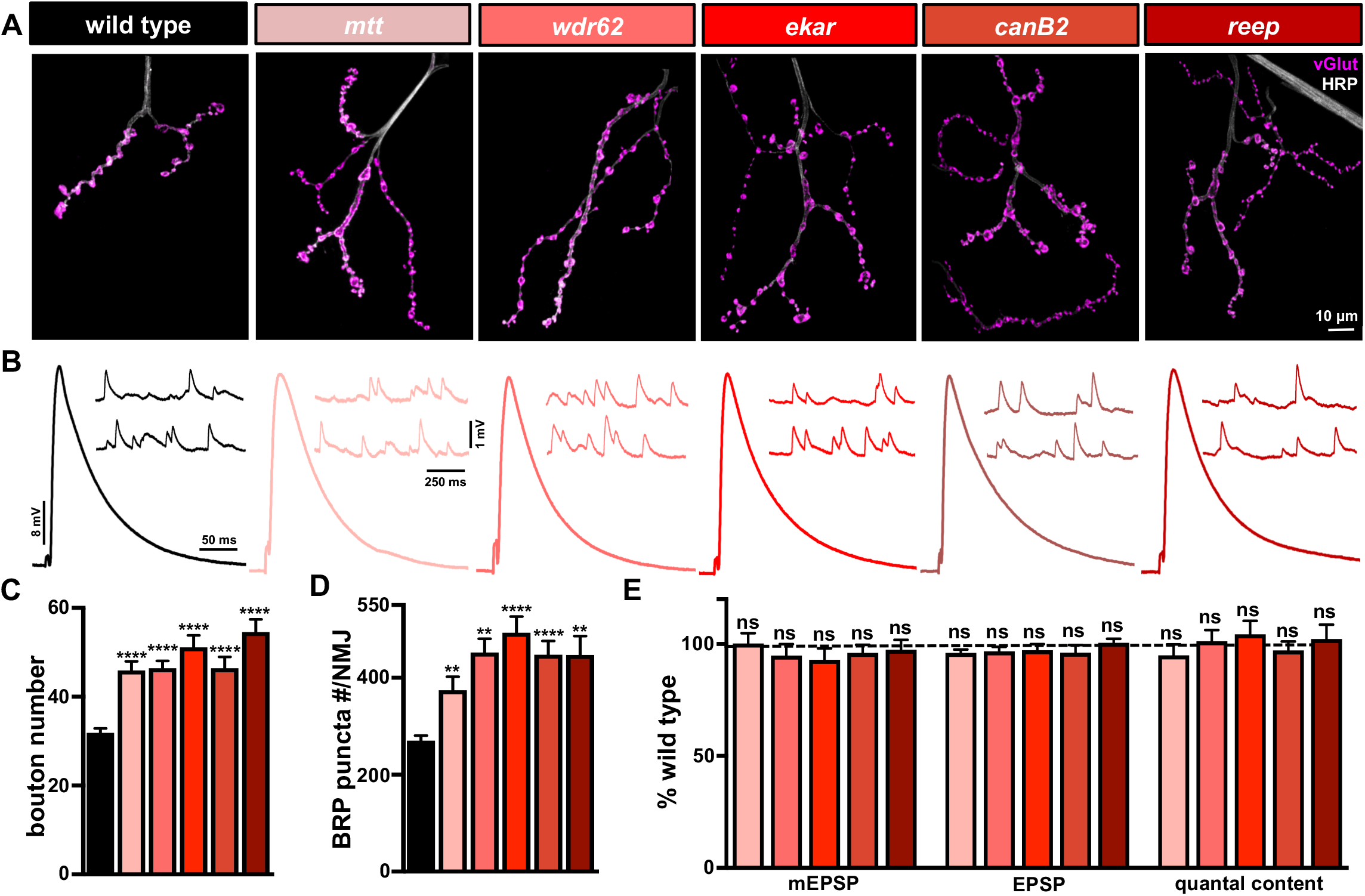
Mutants with enhanced synaptic growth exhibit a concomitant increase in active zone number yet stable levels of synaptic strength. **(A)** Representative images of muscle 4 NMJs in wild type, *mangetout* (*mtt*: *mtt^1^*/*mtt^Df(2R)H3D3^*), *WD repeat domain 62* (*wdr62: wdr62^1^/wdr62^Df(2L)Exel8005^*), *eye-enriched kainate receptor* (*ekar: ekar 1*), *calcineurin B2* (*canB2*: *canB2^1^*/*canB2^Df(2R)BSC265^*), and *receptor expression enhancing protein* (*reep: reep^1^*/*reep^Df(2R)WI345^*) mutants immunostained with anti-HRP and anti-vGlut. **(B)** Representative EPSP and mEPSP traces showing no significant changes in the overgrowth mutants compared to wild type. Quantification of bouton numbers **(C)** and BRP puncta number per NMJ **(D)** in the indicated genotypes reveals a significant increase in both parameters compared to wild type. **(E)** Quantification of mEPSP, EPSP, and quantal content values in the indicated genotypes normalized as a percentage of wild type. Despite enhanced bouton and active zone number per NMJ in the overgrowth mutants, no significant change in presynaptic neurotransmitter release (quantal content) is observed. Error bars indicate ±SEM. One-way analysis of variance (ANOVA) test was performed, followed by a Tukey’s multiple-comparison test. **p≤0.01; ****p≤0.0001; ns=not significant, p>0.05. Detailed statistical information (mean values, SEM, n, p) is shown in S3 Table.

Next, we quantified the total number of BRP puncta per NMJ in these overgrowth mutants. We found an increase in total BRP puncta number per NMJ that correlated with the enhanced synaptic growth observed in each overgrowth mutant (Fig 7A and 7E). Correspondingly, we observed no major differences in bouton size, leading to a parallel increase in total neuronal membrane surface area per NMJ and no change in BRP puncta density (S3 Table). Hence, BRP puncta number essentially scales with bouton number in the overgrowth mutants, in contrast to the undergrowth mutants detailed in Fig 6. This suggests that a reduction in release probability per active zone likely stabilized synaptic strength in these mutants.

The size and abundance of material at individual active zones can vary considerably, and several studies have found that these properties can correlate with release probability [64-66]. At the *Drosophila* NMJ, there is considerable heterogeneity in the size and intensity of the active zone scaffold BRP and other active zone components [67-69]. Furthermore, recent studies have shown that active zones at this NMJ that are endowed with increased intensity and size correlate with increased release probability during baseline transmission and plasticity [17, 70-73]. We therefore considered that while the total number of BRP puncta per NMJ was increased in the overgrowth mutants, there might have been a corresponding change in the area and/or intensity of each puncta that contributed to their modulation of release probability. Analysis of individual BRP puncta revealed a significant reduction in the mean area of BRP puncta in all five synaptic overgrowth mutants (Fig 8A and 8B; S3 Table). Indeed, the average BRP puncta area scaled with total BRP puncta number per NMJ in wild type and in the synaptic overgrowth mutants (Fig 8C; R^2^=0.27, p-value=0.0006). While we did observe a significant inverse correlation (R^2^ value) between BRP puncta number and area, the curve fit of these data points resulted in a lower correlation value, likely due to a narrower distribution. However, the total abundance of BRP per NMJ, reflected in the sum fluorescence intensity of BRP puncta across an entire NMJ, was not significantly different between wild type and the five overgrowth mutants (Fig 8D; S3 Table). Thus, an apparent tuning of active zone size may have compensated for increased number to reduce release probability per active zone and maintain synaptic strength in the overgrowth mutants isolated from the genetic screen.

**Fig 8.**
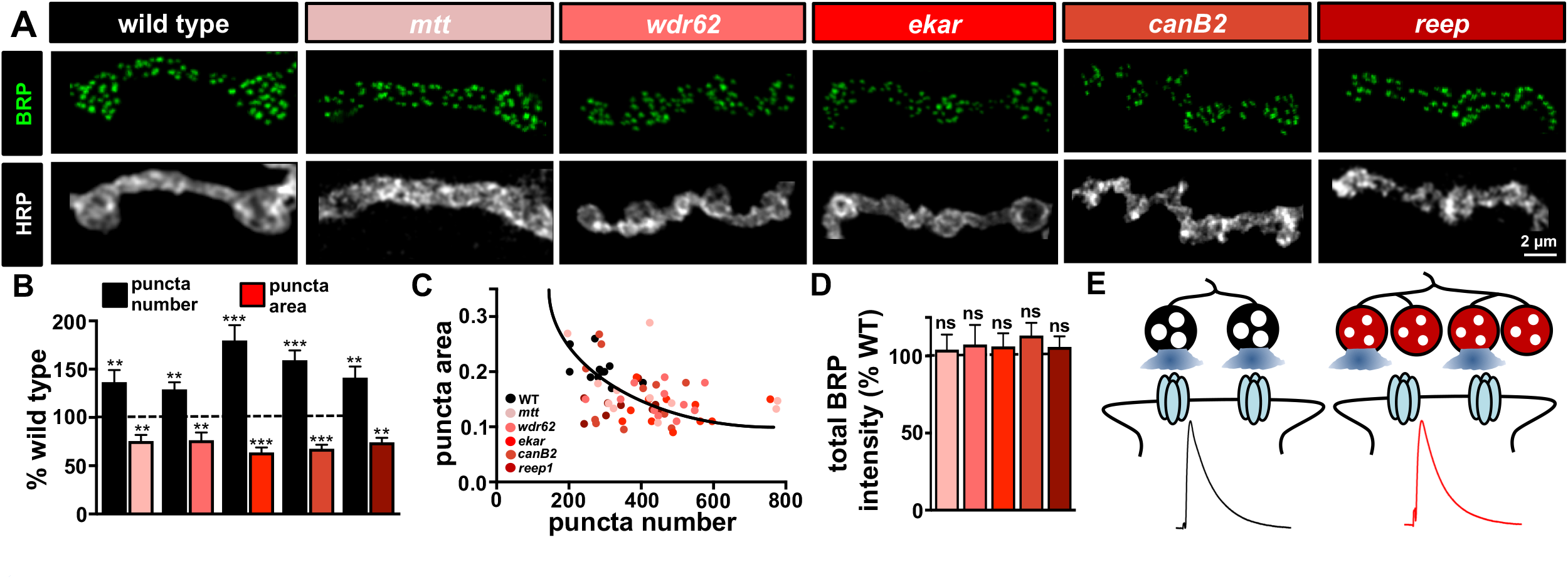
Active zone area is reduced in mutants with enhanced synaptic growth. **(A)** Representative images of individual boutons from wild type and the overgrowth mutants immunostained with anti-BRP and anti-HRP. **(B)** Quantification of BRP puncta number and BRP puncta area in the indicated genotypes normalized to wild-type values. While both bouton and BRP puncta numbers are increased in the overgrowth mutants, a reduction in the average area of each BRP puncta is observed. **(C)** Average BRP puncta area plotted as a function of average BRP puncta number per NMJ in the indicated genotypes demonstrates a homeostatic scaling of BRP puncta area with total number per NMJ, represented by the curve fitted to the data points (R^2^=0.27, p value=0.0006; ***). **(D)** Quantification of total BRP puncta fluorescence intensity per NMJ in the indicated genotypes, suggesting that the total abundance of BRP per NMJ remains unchanged in the overgrowth mutants compared to wild type. **(E)** Schematic illustrating that although both bouton and BRP puncta numbers are increased in overgrowth mutants, a reduction in the area of individual BRP puncta results in reduced release probability per active zone and per bouton to stabilize synaptic strength. Error bars indicate ±SEM. One-way analysis of variance (ANOVA) test was performed, followed by a Tukey’s multiple-comparison test. **p≤0.01; ***p≤0.001; ns=not significant, p>0.05. Detailed statistical information (mean values, SEM, n, p) is shown in S3 Table.

## DISCUSSION

Through a forward genetic screen of ~300 mutants, we have identified genes required for property regulation of synaptic growth and neurotransmission. This approach has revealed several new mutations and RNAi lines that disrupt synaptic growth and function, while also demonstrating that these processes are regulated through distinct pathways. This data implies the existence of a homeostat that stabilizes global synaptic strength while permitting substantial flexibility in synaptic growth. Our analysis has defined three adaptive mechanisms that operate to maintain synaptic strength when synaptic growth is dramatically altered.

### Genes that promote or constrain synaptic growth

A complex repertoire of genes work together to tune synaptic growth, structure, and function. One node of control is the translational modulator FMRP, which has been clearly implicated in the regulation of postsynaptic signaling, dendritic structure, and glutamate receptor dynamics [29, 74-77]. Furthermore, FMRP has also emerged as an important regulator of presynaptic glutamate release via modulation of potassium channels, calcium influx, short-term plasticity, and synaptic vesicle recycling [78-84]. Similarly, genes associated with autism and schizophrenia susceptibility have been shown to have parallel roles in regulating synaptic growth and transmission [85, 86]. Consistent with these studies, our screen identified several disease-linked genes required for proper synaptic growth and transmission. Although further work will be necessary to understand how each gene regulates the growth or function of the synapse, the strength of this large-scale screening approach lies in identifying and assigning functions to individual genes.

There is emerging evidence that both homeostatic and Hebbian forms of plasticity share common genes and signaling networks [8, 87-89]. While the *Drosophila* NMJ is built for stability and has proven to be a powerful model to investigate glutamatergic transmission and homeostatic plasticity, contrasting forms of Hebbian plasticity are less obvious at this synapse. Hence, mutations of genes with specialized functions in non-glutamatergic synaptic transmission or Hebbian plasticity are unlikely to reveal phenotypes using the screening strategy we employed. However, a variety of genes were identified with significant and more subtle roles in regulating synaptic growth and baseline function (S1 Table). Mutations in one gene, *pkc53E*, exhibited reduced synaptic growth and a parallel reduction in transmission, consistent with a scaling model of synaptic growth and transmission. However, our characterization of the remaining synaptic growth mutants revealed evidence for homeostatic adaptations that stabilized synaptic strength across variations in NMJ growth. In the case of the undergrowth mutants *cont* and *Gγ30A*, increased size of individual boutons led to a conservation of both neuronal membrane and active zone number to maintain synaptic strength. Interestingly, there is evidence from studies of other mutants that the size of individual boutons at the *Drosophila* NMJ are inversely correlated with total numbers per NMJ [90-93]. Therefore, adjusting the morphology and size of individual boutons is one adaptive strategy that may generally serve to enable flexibility in synaptic growth while maintaining stable total synapse numbers.

### Homeostatic scaling of glutamate receptor abundance and active zone size

We identified a homeostatic scaling of postsynaptic glutamate receptor abundance that offset reduced presynaptic neurotransmitter release in one synaptic undergrowth mutant. Specifically, *WRNexo* mutants exhibited reduced synaptic growth with a concomitant reduction in presynaptic active zone number and neurotransmitter release. However, this diminished presynaptic efficacy was offset by a compensatory increase in GluRIIA-containing postsynaptic receptors. This phenomenon parallels homeostatic receptor scaling of postsynaptic glutamate receptors following manipulations to activity in mammalian central neurons [94-97]. While glutamate receptors are rapidly and dynamically regulated in central neurons during both Hebbian and homeostatic forms of plasticity [10, 98, 99], receptors at the NMJ are much less dynamic. Glutamate receptors have half lifes of ~24 hr at the *Drosophila* NMJ [100], which parallels the relatively slow dynamics of cholinergic receptors at the mammalian NMJ [101]. However, there is intriguing evidence that postsynaptic receptors at the NMJ can be dynamically regulated in response to changes in presynaptic activity [102, 103], following injury and disease [20, 104-106], and in response to hypo-innervation [20, 21] and similar phenomena occur following injury in the central nervous system [107]. Thus, NMJs may be endowed with an underappreciated degree of latent receptor plasticity mechanisms that can be revealed in response to homeostatic challenges, including synaptic undergrowth.

We identified an apparent homeostatic scaling of active zone size in all five synaptic overgrowth mutants. In contrast to the undergrowth mutants, no changes in bouton size or the postsynaptic sensitivity to neurotransmitter was observed, and active zone number scaled with enhanced synaptic growth. In principle, a variety of compensatory changes in postsynaptic receptors, presynaptic morphology, and/or synapse number could have been homeostatically regulated to maintain synaptic strength. However, all five mutants shared an apparent reduction in the size and intensity of the active zone scaffold BRP, indicative of a functional reduction in release probability of individual active zones. Interestingly, active zone scaffold proteins (CAST/ELKS/BRP) are known to regulate presynaptic release probability by stabilizing calcium channels and the size of the readily releasable synaptic vesicle pool [49, 108-111]. Furthermore, BRP can be rapidly remodeling during homeostatic plasticity to enhance the RRP and promote calcium influx [17, 18, 71, 72, 112]. Finally, a positive correlation between the size and intensity of active zone components and release probability has been observed at the *Drosophila* NMJ [68-70] as well as at vertebrate central synapses [64-66, 113]. Therefore, the reduction in active zone size observed in the overgrowth mutants likely reduces release probability at individual release sites to maintain global NMJ function. More generally, remodeling of active zone structure is an attractive mechanism that might homeostatically tune presynaptic efficacy to stabilize synaptic strength while still permitting flexibility during synaptic growth and pruning.

In the central nervous system, a variety of mechanisms homeostatically scale axonal and dendritic structure and arborization to compensate for altered activity. For example, a homeostatic remodeling of dendritic arborization in the fly visual system is observed in response to chronically elevated or reduced activity [114], and adaptive structural alterations at synapses have been observed during the sleep/wake cycle [4, 115, 116]. Similarly, adaptive changes in the structure and number of dendritic spines are observed in response to imbalances in excitation and inhibition in the central nervous system [2, 117-120]. Parallel adaptations to the axon initial segment and release probability at presynaptic terminals have been demonstrated that counteract homeostatic challenges [65, 89, 121]. Our findings on the interplay between synaptic growth and function underscore the diverse mechanisms that homeostatically stabilize global synaptic strength while permitting dynamic flexibility in the growth of synapses.

## MATERIALS AND METHODS

### Fly Stocks

*Drosophila* stocks were raised at 25°C on standard molasses food. The *w^1118^* strain is used as the wild type control unless otherwise noted, as this is the genetic background of the genetic mutants used in this study. For experiments with the transgenic RNAi lines, control larvae were generated by crossing C15 (*c155-Gal4;Sca-Gal4;BG57-Gal4;* [43]) to *UAS-RFP* (BL 32218). Since the average synaptic growth and electrophysiological values for the mutant control (*w^1118^*) and RNAi control (*c155-Gal4;Sca-Gal4/+;BG57-Gal4/UAS-RFP*) were not significantly different (S1 Table), we pooled all mutant and RNAi line data shown in Figures 1-3. The *WRNexo* null mutants (*WRNexo^**Δ**^*) were previously described [57]. All genetic mutants and transgenic RNAi lines were obtained from the Bloomington Drosophila Stock Center. A complete list of all stocks used in this study, their full genotypes, and their origin can be found in S1 Table.

### Immunocytochemistry

Third-instar larvae were dissected in ice cold 0 Ca^2+^ HL-3 and fixed in Bouin’s fixative for 5 min. Larvae were washed with PBS containing 0.1% Triton X-100 (PBST) for 30 min, and then blocked for an hour with 5% normal donkey serum in PBST. Larvae were incubated overnight in primary antibodies at 4°C followed by a 30 min wash in PBST, 2.5 hour incubation in secondary antibodies at room temperature (20-22°C), a final 30 min wash in PBST, and equilibration in 70% glycerol. Blocking was done with 5% normal donkey serum in PBST. Samples were mounted in VectaShield (Vector Laboratories). The following antibodies were used: mouse anti-Bruchpilot (nc82; 1:100; Developmental Studies Hybridoma Bank; DSHB); rabbit anti-DLG ((1:10,000; [122]); guinea pig anti-vGlut ((1:2000; generated by Cocalico Biologicals using the peptide described in [15]); mouse anti-GluRIIA (8B4D2; 1:100; DSHB); rabbit anti-GluRIIB ((1:1000; generated by Cocalico Biologicals using the peptide described in [59]); guinea pig anti-GluRIID ((1:1000; generated by Cocalico Biologicals using the peptide described in [123]). Donkey anti-mouse, anti-guinea pig, and anti-rabbit Alexa Fluor 488-, Cyanine 3 (Cy3)-, and Dy Light 405- conjugated secondary antibodies (Jackson Immunoresearch) were used at 1:400. Alexa Fluor 647 conjugated goat anti-HRP (Jackson ImmunoResearch) was used at 1:200.

### Imaging and analysis

Samples were imaged using a Nikon A1R Resonant Scanning Confocal microscope equipped with NIS Elements software and a 100x APO 1.4NA oil immersion objective using separate channels with three laser lines (488 nm, 561 nm, and 637 nm). For fluorescence quantifications of BRP intensity levels, z-stacks were obtained using identical settings for all genotypes with z-axis spacing between 0.15 µm to 0.2 µm within an experiment and optimized for detection without saturation of the signal. Boutons were counted using vGlut and HRP-stained NMJ terminals on muscle 6/7 and muscle 4 of segment A3, considering each vGlut puncta to be a bouton. The general analysis toolkit in the NIS Elements software was used for image analysis as described [124]. Neuronal surface area was calculated by creating a mask around the HRP channel that labels the neuronal membrane. BRP puncta number, area, and mean intensity (average intensity of individual BRP puncta) and sum intensity (total intensity of individual BRP puncta) were quantified by applying intensity thresholds and filters to binary layers on the BRP labeled 488 channel. GluRIIA, GluRIIB, and GluRIID puncta intensities were quantified by measuring the total sum intensity of each individual GluR puncta and these values were then averaged per NMJ to get one reading (n). Measurements based on confocal images were taken from at least twelve synapses acquired from at least six different animals.

### Electrophysiology

All dissections and recordings were performed in modified HL-3 saline [125-127] containing (in mM): 70 NaCl, 5 KCl, 10 MgCl_2_, 10 NaHCO_3_, 115 Sucrose, 5 Trehelose, 5 HEPES, and 0.4 CaCl_2_ (unless otherwise specified), pH 7.2. All recordings were performed in 0.4 mM extracellular calcium. Neuromuscular junction sharp electrode (electrode resistance between 10-30 MΩ) recordings were performed on muscles 6 and 7 of abdominal segments A2 and A3 in wandering third-instar larvae. Larvae were dissected and loosely pinned; the guts, trachea, and ventral nerve cord were removed from the larval body walls with the motor nerve cut, and the preparation was perfused several times with HL-3 saline. Recordings were performed on an Olympus BX61 WI microscope using a 40x/0.80 water-dipping objective, and acquired using an Axoclamp 900A amplifier, Digidata 1440A acquisition system and pClamp 10.5 software (Molecular Devices). Electrophysiological sweeps were digitized at 10 kHz and filtered at 1 kHz. Data were analyzed using Clampfit (Molecular devices), MiniAnalysis (Synaptosoft), Excel (Microsoft), and SigmaPlot (Systat) software.

Miniature excitatory postsynaptic potentials (mEPSPs) were recorded in the absence of any stimulation, and cut motor axons were stimulated to elicit excitatory postsynaptic potentials (EPSPs). An ISO-Flex stimulus isolator (A.M.P.I.) was used to modulate the amplitude of stimulatory currents. Intensity was adjusted for each cell, set to consistently elicit responses from both neurons innervating the muscle segment, but avoiding overstimulation. Average mEPSP, EPSP, and quantal content were calculated for each genotype by dividing EPSP amplitude by mEPSP amplitude. Muscle input resistance (R_in_) and resting membrane potential (V_rest_) were monitored during each experiment. Recordings were rejected if the V_rest_ was above -60 mV, if the R_in_ was less than 5 MΩ, or if either measurement deviated by more than 10% during the course of the experiment.

### Experimental Design and Statistical Analysis

For electrophysiological and immunostaining experiments, each NMJ terminal (muscle 6 for physiology, and muscle 4 for immunostaining analyses of synaptic terminals and active zones) is considered an n of 1 since each presynaptic motor neuron terminal is confined to its own muscular hemisegment. For these experiments, muscles 4 or 6 were analyzed from hemisegments A3 for each larvae, and thus each larvae contributes 2 NMJs per experiment. To control for variability between larvae within a genotype, for immunostaining experiments involving BRP and GluRIII, NMJs were analyzed from no less than 6 individual larvae.

Statistical analysis was performed using GraphPad Prism software. Data were tested for normality using a D’Agostino-Pearson omnibus normality test. Normally distributed data were analyzed for statistical significance using a t-test (pairwise comparison), or an analysis of variance (ANOVA) and Tukey’s test for multiple comparisons. For non-normally distributed data, Wilcoxon rank-sum test or Dunn’s multiple comparisons after nonparametric ANOVA were used. All data are presented as mean +/-SEM. with varying levels of significance assessed as p<0.05 (*), p<0.01 (**), p<0.001 (***), p<0.0001 (****), ns=not significant. See S3 Table for additional statistical details and values.

## ACKNOWLEDGEMENTS

We thank the Dr. Mitch McVey (Department of Biology, Tufts University, USA) for sharing *WRNexo* mutants. We acknowledge the Developmental Studies Hybridoma Bank (Iowa, USA) for antibodies used in this study, and the Bloomington Drosophila Stock Center (NIH P4OD018537) for fly stocks. We also acknowledge Andrew Tung, Andrew An, Luke Nunnely, and Michelle Chee for contributions during early phases of this project. MK and SH were supported by USC Undergraduate Research Awards, and PG and KK were supported in part by USC Provost Graduate Research Fellowships. This work was supported by a grant from the National Institutes of Health (NS091546) to DD.

## AUTHOR CONTRIBUTIONS

PG, MK, and SH obtained all experimental data. BK and KK organized and supervised early phases of the FMRP screen. MK and PG analyzed all data. The manuscript was written by PG and DD with comments from MK.

---

**S1 Table. Quantification of synaptic growth and function in all genes, mutants, and RNAi lines screened.** The Flybase ID, CG number, gene name, putative function, full mutant or RNAi genotype, and source (BDSC stock number) for each fly stock screened is noted. Further, quantification of bouton number, mEPSP amplitude, EPSP amplitude, and quantal content for each line is shown.

**S2 Table. Putative functions of synaptic undergrowth and overgrowth genes.** Putative functions of each synaptic undergrowth and overgrowth gene is shown along with related references.

**S3 Table. Absolute values for normalized data and additional statistics.** The figure and panel, genotype, and experimental conditions are noted. For electrophysiological recordings, average mEPSP, EPSP, quantal content (QC), resting potential, input resistance, number of data samples (n), p values, and significance values are shown, with standard error noted in parentheses. For analysis of confocal images, average fluorescence intensity values and related parameters are shown. Standard error values are noted in parentheses. Rows highlighted in blue are the respective controls or baseline values for the particular experiment being referenced.

